# Ubi-SCAPE enables deep exploration of the poly-ubiquitylome

**DOI:** 10.64898/2026.02.13.703496

**Authors:** Harvey E. Johnston, Andrew Frey, Gaurav Barve, Sam Carling, Matthias Trost, Rahul S. Samant

## Abstract

Post-translational modification with chains of the 76-amino-acid protein ubiquitin (‘poly-ubiquitylation’) confers diverse fates to the targeted protein and non-protein substrates, including degradation, intracellular trafficking, and signal transduction. Despite being one of the most frequent modifications, the complexity of poly-ubiquitin chains adds methodological challenges to their characterization. Trypsin-resistant tandem-ubiquitin-binding entities (trTUBEs), engineered from natural ubiquitin-binding domains, can capture intact poly-ubiquitylated proteins with high cumulative avidity. However, such approaches have suffered from considerable co-eluting contaminants in proteomics applications. Here, we introduce an optimized trTUBE-based method for poly-ubiquitylated proteome purification, drastically depleting non-ubiquitylated protein contaminants and mono-ubiquitylated proteins. The method, termed Ubiquitylomics by Stringent, Cleavable, Affinity-based Proteome Extraction (Ubi-SCAPE), offers a streamlined and reproducible (median R^2^ > 0.98; CV < 8%) means of characterizing the poly-ubiquitylome. Over 7,800 poly-ubiquitylated proteins and 8,500 ubiquitin-modified peptides (diGly) were quantified with Ubi-SCAPE at a throughput of 40 samples per day, as well as over 6,000 from an equivalent of 33 μg unstressed cell lysate. Upon acute stress by heat-shock, we identified 2,700 proteins and 8,000 diGly peptides with increased poly-ubiquitylation—offering similar biological insight as with far more material-, labor-, and cost-intensive peptide-based ubiquitin enrichment methods. Ubi-SCAPE therefore provides a simple and effective means of comprehensively quantifying a selective-enriched poly-ubiquitylome.

**GRAPHICAL ABSTRACT:** 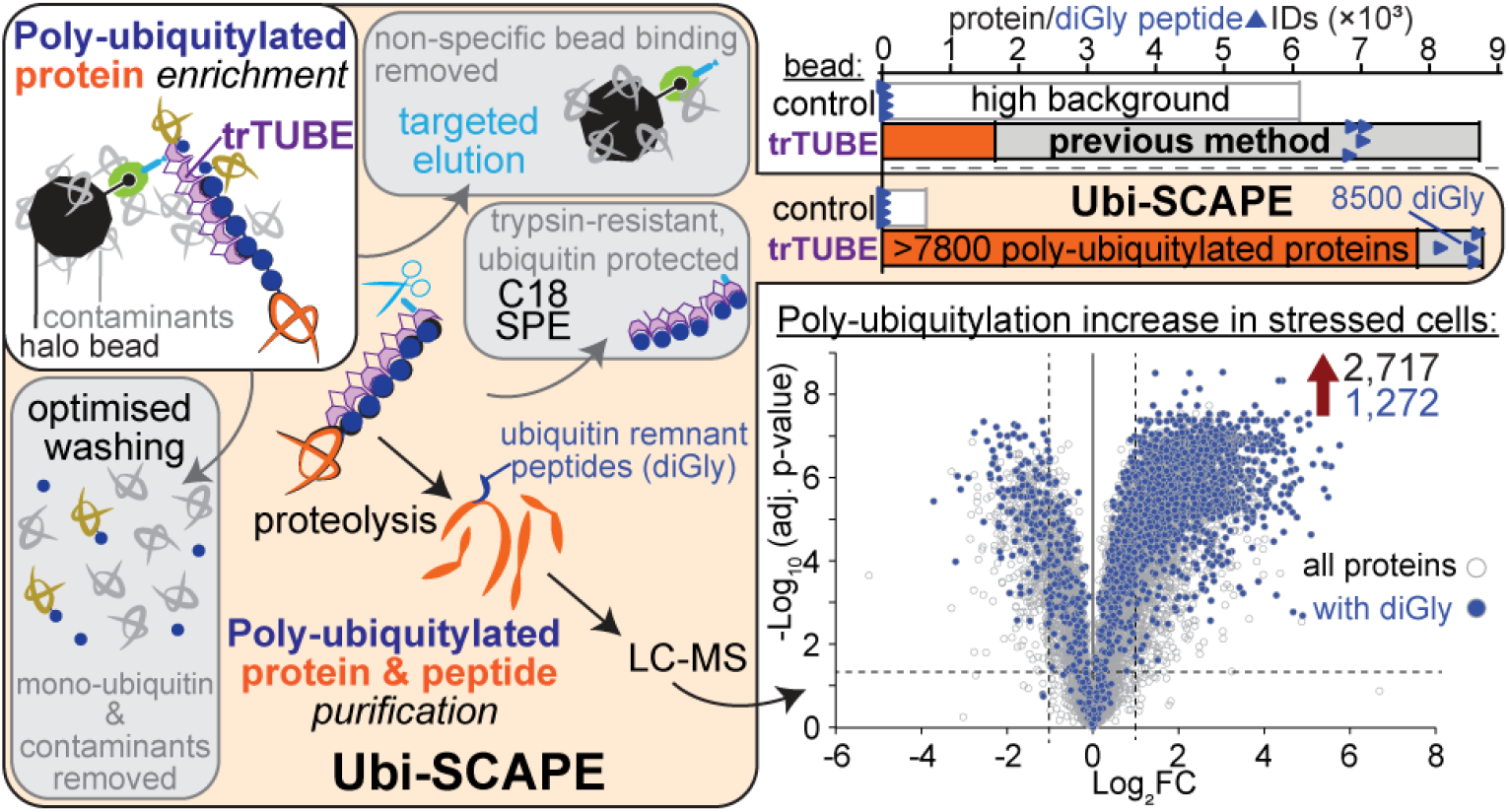

## INTRODUCTION

Ubiquitin is a highly-conserved and abundant post-translational modifier throughout the proteome of eukaryotic organisms.^1^ Although post-translational modification (PTM) of substrates with one or more ubiquitin units is most commonly associated with targeting them for degradation, ‘ubiquitylation’ has highly diverse roles, and potentially represents the most frequent posttranslational modification in the human proteome.^2, 3^ Comprehensive identification and quantification of ubiquitylated proteins therefore promises substantial insight into cellular pathways and states such as protein misfolding, degradation and signalling.^4, 5^ Rather than representing a uniform modification, ubiquitylation commonly exists as a range of mono-, multi-mono, or poly-ubiquitin chains on the original protein substrate. Like other critical PTMs, the conservation and frequency of ubiquitylation across eukaryotes reinforces a universal role across biological processes. Despite this biological importance, tools for its characterization remain relatively limited and infrequently applied.

Current gold-standard approaches to quantify the ubiquitin-modified proteome (‘ubiquitylomics’) involves enrichment of peptides bearing the “diGly” ubiquitin-remnant fingerprint after tryptic digestion, using proprietary antibodies raised against that motif.^2, 6-9^ Although this approach is effective for isolating such peptides, it is resource-intensive, requires large amounts of starting material and multiple specialized steps, and cannot distinguish between mono-, poly- and some ubiquitin-like modifications.^6^ Other tools for the isolation of all free ubiquitin and ubiquitylated proteins have also been developed;^10-18^ the complexity and diversity of poly-ubiquitylation, however, has remained challenging to characterize.

TUBEs (Tandem-Ubiquitin-Binding-Entities) were developed for this purpose,^19^ employing a string of ubiquitin-binding domains to capture ubiquitin chains through high avidity interactions. Trypsin-resistant TUBEs (trTUBEs) represented an additional improvement, protecting both the TUBE and the bound ubiquitin from tryptic digestion during bottom-up proteomics workflows, thereby reducing the signal from these two major “contaminants” and improving depth of coverage for the ubiquitin-modified proteome itself.^11^

Although TUBEs and trTUBEs have been employed in proteomics applications,^11, 20-24^ their uptake for poly-ubiquitylomics has been limited, perhaps due to the risk of contamination from abundant or co-precipitating proteins that were not themselves poly-ubiquitylated—a challenge for many high-sensitivity proteomics affinity purification workflows.^25^

Here, we present Ubi-SCAPE (Ubiquitylomics by Stringent, Cleavable, Affinity-based Proteome Extraction), a trTUBE-based poly-ubiquitylomics pipeline that yields unprecedented coverage, reproducibility, and specificity, and provides clear improvements over existing ubiquitin-enrichment approaches.

## MATERIALS AND METHODS

### Materials and reagents

All key materials and reagents are listed in the Key Resources Table.

### Construction of pET28aOPT HaloTag–TEVc–empty and HaloTag–TEVc–6×trTUBE

To construct pET28aOPT HaloTag-TEVc-empty, a fragment containing the self-labelling HaloTag, the Tobacco Etch Virus protease (TEVp) cleavage site ENFLQG (denoted ‘TEVc’ here), and a multiple-cloning-site (MCS), from pET28a HaloTag-TEVc-[MCS] (DU23222, MRC-PPU Reagents & Services) was PCR amplified with custom primers 108-BlpI-F1 (ATTATGCTTAGCGGCAGCAGCCATC) and pET28a-BlpI-R1 (TGCTAGTTATTGCTCAGCGGTGGCA), followed by BlpI digest and ligation into the optimised pET28a T7pCONS TIR-2 sfGFP (gift from Daniel Daley, Addgene #154464).^26^ To construct pET28aOPT HaloTag-TEVc-6×trTUBE, the 6×trTUBE insert from pRSET-6×TR-TUBE (gift from Yasushi Saeki, Addgene #110313)^18^ was double-digested with EcoRV + DraIII and ligated into Eco53kI + DraIII double-digested pET28aOPT HaloTag-TEVc-empty. To generate an alternative version of the construct with an additional 6×His tag followed by a thrombin cleavage site for affinity purification (‘pET28aOPT 6×His-HaloTag-TEVc-6×trTUBE’), the 6×His-HaloTag fragment from pET28a 6×His-HaloTag-Nrf2 (gift from Yimon Aye, Addgene #62455) was double-digested with XhoI + XbaI and ligated into similarly-digested pET28aOPT HaloTag-TEVc-6×trTUBE. pET28aOPT 6×His-HaloTag-TEVc-4×trTUBE and pET28aOPT 6×His-HaloTag-TEVc-4×trTUBE_mut_ were constructed by DraIII + BglII sequential digest of pET28a 6×His-TEVc-HaloTag-4×trTUBE (MRC-PPU Reagents & Services, #DU58810) or pET28a 6×His-TEVc-HaloTag-4×trTUBE_mut_ (MRC-PPU Reagents & Services, #DU58829)(both gifts from Yu-Chiang Lai, U. Birmingham) and ligation into similarly-digested pET28aOPT 6×His-HaloTag-TEVc-6×trTUBE.

All cloning and propagation was performed in sub-cloning-efficiency competent DH5-alpha *E. coli* (Thermo Fisher Scientific; Cat# 18265017), and all enzymes, buffers, and kits for cloning were from New England Biolabs. All plasmids were verified by whole-plasmid sequencing (Plasmidsaurus).

### trTUBE expression and conjugation to HaloTag magnetic beads

Rossetta 2(DE3) competent *E. coli* (Novagen/Sigma; Cat# 71397) transformed with one of pET28aOPT HaloTag–TEVc–6×trTUBE, pET28aOPT HaloTag–TEVc–empty, pET28aOPT 6×His-HaloTag-TEVc-4×trTUBE, or pET28aOPT 6×His-HaloTag-TEVc-4×trTUBE_mut_ plasmids were cultured at 37 °C for 18 h to OD_600_ ∼0.7, adjusted to 1 mM IPTG (Melford Laboratories, Cat# B1008) to induce expression, and incubated for 24 h at 16 °C. *E. coli* were pelleted at 7,000 ×*g* for 10 min at 4 °C. After supernatant aspiration, bacteria were lysed with three pellet volumes (w/v) of bacterial lysis buffer (50 mM Tris HCl, pH 7.5, 150 mM NaCl, 1% Triton X 100, 20 mM TCEP, 1× cOmplete protease inhibitors, 0.2 mM phenylmethylsulfonyl fluoride (PMSF)) with sonication on ice for 2 min total at 20 W (10 s intervals) using a non-microtip probe (Covaris). Lysates were cleared by centrifugation for 30 min at 16,000 ×*g* at 4 °C, and the supernatant snap-frozen in liquid nitrogen and stored at –70 °C as 200 μL aliquots.

Magne-HaloTag Beads (Promega, Cat# G7281) saturation with HaloTag–TEVc–6×trTUBE protein was determined (Fig. S1) as approximately 140 μL *E. coli* lysate as prepared above per 1 mL of 20% Magne-HaloTag beads. To accommodate for batch variability, 200 μL *E. coli* lysate was used per 1 mL for all preparations. Magne-HaloTag beads were washed 3 times with 1 mL of IP buffer (50 mM Tris-HCl, pH 7.5, 150 mM NaCl, 1% IGEPAL CA-630), and incubated with 200 μL *E. coli* lysate for 2 h at room temperature with end-over-end rotation at 20 rpm. Beads were then washed three times with 1 mL of trTUBE wash buffer (defined below), three times with IP buffer, and stored at 4 °C as a 2% slurry.

### Cell culture and heat-shock

HEK293 cells were grown in DMEM (Gibco; Cat# 41965039) supplemented with 10% heat-inactivated HyClone Characterized Fetal Bovine Serum (FBS)(Gibco; Cat# SH30071.02HI), 100 U/mL penicillin-streptomycin (Gibco; Cat# 15140122), 2 mM L-glutamine (Gibco; Cat# 25030081), and 1× non-essential amino-acids (NEAA)(Gibco; Cat# 11140050), in a humidified incubator set at 37 °C with 5% CO_2_. Cells were plated from a single pool of cells and grown to ∼80% confluency and were either left at 37 °C (‘unstressed’) or incubated at 44 °C for 2 h (‘heat-shock’). Cells were aspirated, resuspended in 10 mL of ice-cold phosphate-buffered saline (PBS)(Sigma; Cat# P2272) per 15 cm plate, and washed three times with 10 mL PBS per 15 cm plate, pelleted at 300 ×*g*, fully aspirated, and snap-frozen in liquid nitrogen.

### Cell lysate preparation

Cell pellets were lysed on ice in 4 cell-pellet volumes of chilled, modified SP3 buffer,^27^ termed ‘urea SP3 lysis buffer’ (50 mM HEPES pH 8.0, 8 M urea, 1% SDS, 1% Triton X-100, 1% IGEPAL CA-630, 1% Tween 20, 1% sodium deoxycholate, 150 mM NaCl, 10 mM TCEP, 1x cOmplete protease inhibitors, EDTA-free, 1 mM PMSF, 10 µM PR-619, 40 mM 2-chloroacetamide (CAA), 20 U/μL Pierce Universal Nucleases (Thermo; Cat# 88701)). Lysates were incubated on ice for 15 min and transferred to a thermomixer for 30 min at 25 °C, 800 rpm for reduction, alkylation, and nuclease action. Lysates were cleared at 16,000 ×*g* for 5 min at 4 °C. Protein concentration was estimated by BCA assay (Thermo; Cat# 23225) of peptides generated by SP4 protein clean-up,^28^ and lysates adjusted to 5 µg/µL.

### Poly-ubiquitylome purification with trTUBE

Affinity purifications were performed with 500 μg of BCA-quantified cell lysate (or 100 μg, where specified), typically in 100 µL of lysis buffer, diluted in 1,700 µL of IP buffer, with 200 µL of 2% HaloTag bead:IP buffer slurry to give a final lysate:IP buffer dilution of 1:20, and incubated for 18 h at 4 °C with end-over-end rotation at 20 rpm. Replicates were defined as full technical repeats of independent lysate:bead incubations. Beads were gathered by a magnet, aspirated, and washed three times (each with 30× manual end-over-end inversions over ∼1 min) with 1 mL of either a ‘typical’ wash buffer (IP buffer adjusted to 500 mM NaCl)—representative of wash buffers previously used for TUBE enrichments^11, 20-24^ —or an optimized ‘trTUBE wash buffer’ (IP buffer supplemented with 5% SP3 buffer, 2 M urea, 1 M NaCl). Beads were subsequently washed three times with 1 mL 50 mM HEPES pH 8.0. For the final wash, beads were transferred to a new 1.5 mL Protein LoBind tube (Eppendorf) and fully aspirated with the aid of brief centrifugation (2,000 ×*g*, 15 s). Tubes were placed on ice and 1 µL (10 U) of Tobacco Etch Virus protease (TEVp)(New England Biolabs, MA USA; Cat# P8112) in 20 µL 50 mM HEPES pH 8.0 was added and incubated for 5 min prior to transferring to a thermomixer for 2 h at 30 °C, 1,000 rpm to elute poly-ubiquitylated proteins from the beads. Beads were gathered by magnet, with the supernatant containing complexes of TEVp-eluted, trTUBE-bound poly-ubiquitylated proteins.

### Recombinant poly-ubiquitin capture with trTUBE

100 ng of each human recombinant di-ubiquitin chain (K6-, K11-, K29-, K33-, K48-, and K63-linked) (South Bay Bio; SBB-UP0060, 0063, 0077, 0066, 0069, and 0072, respectively), tetra-ubiquitin chain (K6-, K11-, K29-, K33-, K48-, and K63-linked)(South Bay Bio; SBB-UP0061, 0064, 0078, 0067, 0070, and 0073, respectively), linear M1-linked poly-ubiquitin chains (Ub2– 7) (Enzo; BML-UW1010), and mono-ubiquitin (Sigma; Cat# U5507) were incubated with trTUBE as described above. Elutions were performed either sequentially, as detailed in each figure, using 15 µL of each potential elution condition for 2 minutes at 4 °C and 800 rpm, or in 1× SDS–PAGE loading buffer (25 mM Tris-HCl pH 6.8, 10% glycerol, 1% SDS, 0.05% (w/v) Orange G) for 10 min at 75 °C and 800 rpm in a thermomixer.

### SDS–PAGE, staining, and immunoblotting

Protein solutions in 1× SDS–PAGE loading buffer were run on 10% or 15% Tris-Glycine-SDS polyacrylamide gels, and stained with Coomassie dye for total protein, or with mouse anti-HaloTag (Promega; Cat# G9211) or mouse anti-ubiquitin (LifeSensors, Cat# VU101) primary antibody followed by either DyLight 680 or DyLight 800-conjugated anti-mouse IgG secondary antibody (Cell Signaling Technology; Cat# 5470 or 5257, respectively). Immunoblots were visualized on a LiCor Odyssey CLx Imager (LI-COR Biotech).

### Protein digestion

TEVp-eluted proteins were added to Protein LoBind tubes (Eppendorf) containing 50 ng of trypsin and Lys-C (Promega; Cat# V5073) in 10 µL of 50 mM HEPES pH 8.0 (prepared as a master-mix with tubes incubated on ice for 5 min). For on-bead digests, 10 µL of the above master-mix was added to the aspirated beads suspended in 20 µL of 50 mM HEPES pH 8.0. Digests were incubated for 2 h at 47 °C on a thermomixer (Eppendorf) at 1000 rpm. Post-digestion, samples were acidified to a final concentration of 1% LC-MS-grade formic acid (Fisher).

### LC–MS proteomics

Liquid chromatography (LC) was performed using an Evosep One system with a 15 cm Aurora Elite C18 column with integrated CaptiveSpray emitter (IonOpticks), at 50 °C. Buffer A was 0.1% formic acid in HPLC water, buffer B was 0.1% formic acid in acetonitrile. Immediately prior to LC-MS, 10 µL (one-third) of the resulting peptides were loaded onto the LC system– specific C18 EvoTips, according to manufacturer instructions, and subjected to the predefined WhisperZoom 40 SPD protocol (where the gradient is 0–35 % buffer B, 200 nL/min, for 32.5 min, 40 SPD are permitted using this method). The Evosep One was used in line with a timsToF-HT mass spectrometer (Bruker), operated in diaPASEF mode. Mass and IM ranges at tims-MS1 were 100–1700 m/z and 0.75–1.45 1/K0. MS2 during diaPASEF was performed using 16 variable width IM-m/z windows across two PASEF ramps (Table S3) designed using py_diAID^29^ as part of an ongoing ubiquitylation study. Tims accumulation and ramp times were 100 ms with a ramp rate of 9.42 Hz, total cycle time was ∼1.8 s. Collision energy was applied in a linear fashion, where ion mobility = 0.6–1.6 1/K0, and collision energy = 20–59 eV.

### LC-MS data processing

Raw data were initially processed by DIA-NN (v.1.9.0) using its *in silico* generated spectral library function.^30^ Reference proteome FASTA files included *H. sapiens* (UP000001414, downloaded from UniProt on 23/01/2023) and *E. coli* (UP000000625, downloaded from UniProt on 25/04/2023), and a common contaminants list with the addition of TEVp and the HaloTag sequences.^25^ In addition, the RPS27A and UBA52 protein reference sequences were truncated to exclude the ubiquitin fusion open-reading-frame, and the polyubiquitin proteins (UBB and UBC) were truncated to di-ubiquitin for ease of annotation. Trypsin specificity with a maximum of 2 variable modifications and 1 missed cleavage were permitted per peptide; cysteine carbamidomethylation was set as a fixed modification; and oxidation of methionine and K-GG (diGly-lysine, missed cleavage required) were set as variable modifications. Peptide length and m/z was 7–30 and 300–1256, and charge states 2–4 were included. Mass accuracy was fixed to 15 ppm for both MS1 and MS2. Protein and peptide FDR were both set to 1%. Match between runs was used. Normalisation was disabled to precisely assess technical effects. All other settings were left as defaults.

### Data analysis and bioinformatics

Proteomics data and diGly remnant peptides are detailed in Tables S1 & S2. Proteins were strictly defined as a contaminant if they were observed for any of the 4 non-binding bead replicates (HaloTag–TEVc–empty) for a respective condition and subtracted from the list defined as ‘confident’ identifications. Proteins reaching an FDR-adjusted p-value of p < 0.05 and a fold-change of at least 2-fold upon heat-shock (absolute Log_2_FC > 1) were considered to significantly over-abundant poly-ubiquitylated proteins. Gene Ontology (GO) term enrichment analysis and functional annotation enrichment was performed with Database for Annotation, Visualization, and Integrated Discovery (DAVID)(v2023q4).^31^ Terms were filtered to include those with Benjamini-adjusted significance of p < 0.05. StringDB (v11)^32^ was used to visualise the top 200 significantly increased poly-ubiquitylated proteins in heat-shock, which was trimmed of unconnected proteins, confidence set to > 0.7, with the specified terms highlighted for proteins annotated, selected based on both term-enrichment significance and number of proteins.^32^ A custom R script utilising the diann-R package and R (v4.2.2) was used to generate MaxLFQ-based intensity tables for protein groups, precursors, and diGly peptides from the main report file (.parquet) generated by DIA-NN. Although the majority of diGly residues were internal, a minority were annotated as C-terminal. On inspection of these peptides, most of these (823/1,010) contained an additional uncleaved lysine residue within 5 amino acids of the C-terminally annotated diGly site, none of which were observed in any non-binding control samples; therefore, these were considered to be misannotated and were included in analyses. The MS proteomics data have been deposited to the ProteomeXchange Consortium (http://proteomecentral.proteomexchange.org) via the PRIDE partner repository with the data set identifier PXD073597.^33^

## RESULTS AND DISCUSSION

### Establishing a stringent poly-ubiquitylated proteome enrichment pipeline

Like many affinity purifications, TUBE-based enrichment of poly-ubiquitylated proteins from complex, high dynamic-range protein mixtures present a challenge in distinguishing true substrates from non-specific binding or contamination from abundant or adhesive proteins.^34^ Existing applications have employed relatively cautious binding and washing conditions, with non-ionic detergents and salts as the only means of minimizing native protein:protein and protein:bead interactions^11, 20-24^. By contrast, final elution of bound substrates often employ harsh denaturants, such as strong ionic detergent, heat, or on-bead trypsin proteolysis. We reasoned that the combination of these two factors would compound the presence of highly-abundant and adhesive proteins in the final eluate analyzed by LC-MS, and thus substantially limit the confidence of true-positives observed by TUBE-based poly-ubiquitylomics.

To reduce these concerns, we designed a workflow for poly-ubiquitylated proteome purification via an engineered HaloTag-fused 6×trTUBE construct (i.e., six tandem repeats of the ubiquitin-associated domain from human UBQLN1, with all Lys and Arg residues mutated to Ala)^11^, with a Tobacco Etch Virus protease (TEVp) cleavage site introduced in between the HaloTag and the 6×trTUBE that would offer a specific, targeted means of elution (Fig. 1A–B). We next performed a set of experiments to define the ideal binding conditions, including magnetic-HaloTag bead saturation, stringent washing conditions, and elution conditions (Figs. S1–S3). The optimized protocol employed harsh initial lysis to denature protein and dissociate interactors, followed by the minimal dilution required to restore trTUBE:poly-ubiquitin binding. Similar principles were applied to the wash conditions: we defined the harshest buffer conditions that maintained trTUBE:poly-ubiquitin binding, to ensure the lowest possible contamination from non-specific interactors. Building upon earlier evidence that trTUBE binds longer poly-ubiquitin chains better than shorter chains,^18^ we found that the denaturant tolerance of the poly-ubiquitin:trTUBE interaction was proportional to chain length (Fig. 1C). This observation allowed us to select binding and wash conditions that minimized retention of mono- and di-ubiquitin. Harsher washing conditions promised a greater depletion of high-abundance proteins, such as ribosomes and histones, while maintaining poly-ubiquitin binding. In addition to their high-abundance, histone proteins in particular are canonically mono-ubiquitylated (as an epigenetic mark);^35^ therefore, depletion of histones and other abundant mono-ubiquitylated proteins would result in an increased depth-of-coverage into other, less-abundant substrates.

**Figure 1.**
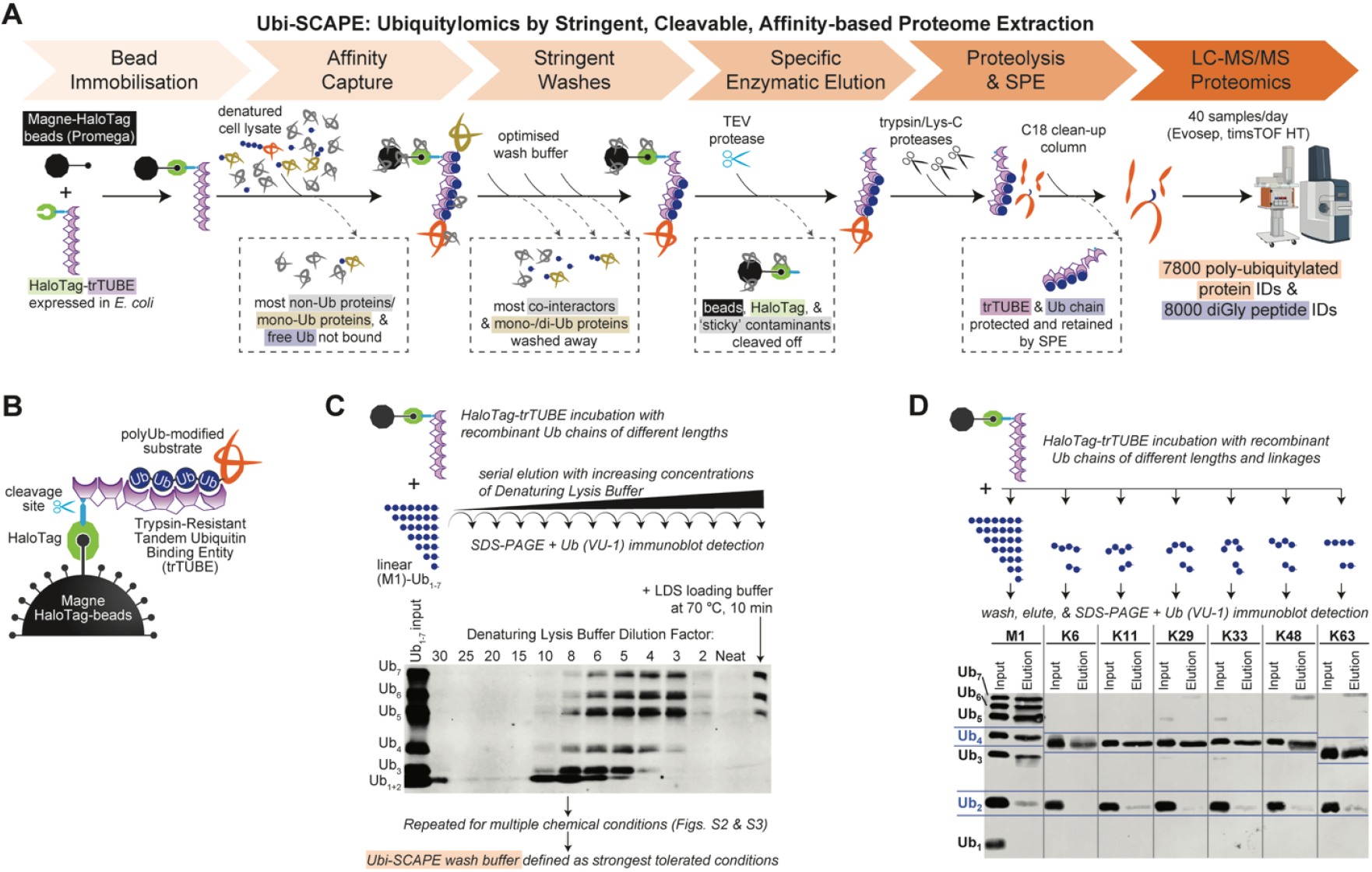
Establishment of the Ubi-SCAPE pipeline for high purity poly-ubiquitin chain purification. (A) Ubi-SCAPE workflow for purification and quantification of poly-ubiquitylated proteins from diverse biological material, with stringent washes, elution by targeted release, and subsequent characterisation by LC-MS-based proteomics of purified, trypsin-digested poly-ubiquitylated protein peptides. (B) A trypsin-resistant tandem-ubiquitin-binding-entity construct (HaloTag–TEVc–6×trTUBE) conjugated to Magne-HaloTag beads binding to poly-ubiquitylated proteins. (C) Poly-ubiquitin chain length is proportional to detergent-resistance of 6×trTUBE binding, tested using linear (M1-linked) poly-ubiquitin of various lengths (1–7) and iterative incubations to quantify dissociation. (D) trTUBE-based affinity capture is poly-ubiquitin linkage agnostic, excluding mono- and the vast majority of di-ubiquitin.

The surface area of beads was also a suspected source of contamination, where harsh elution mechanisms could co-dissociate non-specifically-bound proteins. Targeted elution of the trTUBE and bound substrate away from the HaloTag-beads via TEVp cleavage prior to trypsin digestion presented a promising means of reducing contaminating proteins, in comparison with direct on-bead trypsin digestion.

Our optimized conditions also maintained the linkage non-selective nature of trTUBE (Fig. 1D), thus allowing unbiased poly-ubiquitin chain capture.

By employing stringent washing, our *in vitro* work indicated that Ubi-SCAPE provided more confident purification of the poly-ubiquitylome than previous TUBE-based methods.

### Ubi-SCAPE purifies the poly-ubiquitylome

To evaluate the performance of Ubi-SCAPE vs. the most commonly adopted published TUBE-based protocols, a comparative poly-ubiquitylome purification was conducted comparing HaloTag–TEVc–6×trTUBE with a ‘non-binding’ empty HaloTag-only construct (HaloTag– TEVc–empty) to define entirely non-specific protein contamination (Fig. 2A). We also evaluated previously described variants of trTUBE with fewer tandem domains, or lower binding capacities, for poly-ubiquitin capture (discussed later). Lysates from HEK293 cells in unstressed or heat-shocked conditions were used to evaluate performance in physiologically-relevant basal or elevated poly-ubiquitylation settings, respectively. In addition to comparing the trTUBE variants, we also compared our stringent wash conditions with the more typical wash conditions employed previously.^22^

**Figure 2.**
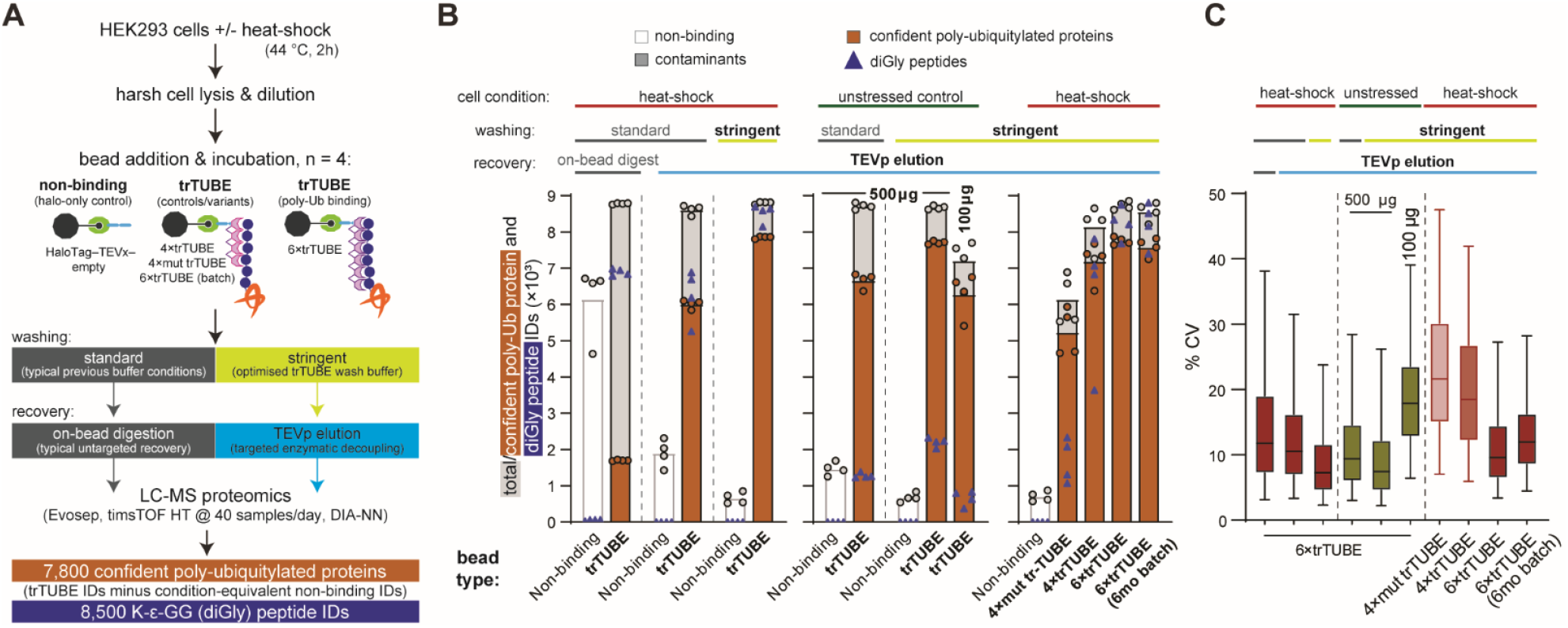
Ubi-SCAPE reproducibly purifies the poly-ubiquitylome with negligible non-specific binding. (A) The Ubi-SCAPE method, incorporating stringent washing and targeted elution, was compared with existing TUBE-based methods employing the equivalent of on-bead digest elution and low-detergent, high-salt washing. For each comparison, trTUBE (HaloTag– TEVc–6×trTUBE) and non-binding (HaloTag–TEVc–empty) bead incubations were performed for 500 μg of lysate, to evaluate poly-ubiquitin capture and the extent of non-specific binding. Both stressed (heat-shock for 2 h at 44 °C) and unstressed cells were evaluated, alongside a more limited-input 100 μg test. Bead variants were also compared, including a binding-deficient mutated HaloTag–TEVc–4×trTUBE_mut_, a shorter HaloTag–TEVc–4×trTUBE, and a batch-stability control for HaloTag–TEVc–6×trTUBE beads stored for 6 months at 4 °C. (B) LC-MS proteomics identifications for evaluations detailed in (A). Confident poly-ubiquitylated proteins (orange) are those IDs observed amongst total IDs (grey) that were not identified in the corresponding non-binding HaloTag–TEVc–empty (white, deemed as contaminants). Lys-ε-GlyGly (diGly) ubiquitin remnant peptide IDs are also shown (blue triangles). (C) Percentage coefficients of variation (% CV) for those confident poly-ubiquitylated proteins IDs defined in (B).

We also employed stringent criteria for assigning confident ‘true positives’, with only proteins not observed to elute from non-binding beads (HaloTag–TEVc–empty) in any of the four replicates as being assigned as confident poly-ubiquitylated proteins (Fig. 2B). Although there were broadly similar numbers of total protein identifications with the HaloTag–TEVc– 6×trTUBE across our tested wash and elution parameters, there was a clear difference in terms of number of proteins identified with the non-binding beads (assigned as contaminants), which had a knock-on effect on the number of poly-ubiquitylated proteins that could be confidently assigned.

As hypothesised, TEVp elution consistently yielded fewer contaminants than on-bead digestion. With standard (non-stringent) washes, we identified up to 6,716 vs. 2,315 non-binding-bead-derived contaminant protein IDs, and therefore 1,700 vs. 6,700 confident poly-ubiquitylated proteins, for on-bead digestion vs. TEVp elution, respectively. However, standard wash conditions exhibited relatively low poly-ubiquitylated protein reproducibility (Fig. 2C) and protein intensities that correlated with the input lysate proteome (R^2^ = 0.5 and 0.23 for on-bead digestion and TEVp elution, respectively), suggesting co-purification of high-abundance contaminants (Fig. S4A). By contrast, the Ubi-SCAPE protocol, employing both stringent washing and TEVp-assisted targeted enzymatic release, provided over 7,800 confident poly-ubiquitylated proteins, reduced contamination to fewer than 850 proteins, exhibited negligible correlation with the input proteome (R^2^ = 0.046; Fig. S4A), and displayed the highest reproducibility (median R^2^ > 0.98, CV < 8%; Fig. 2C). Additionally, over 8,500 diGly ubiquitin remnant peptides, mapping to over 2,500 proteins, were identified for cells stressed with heat-shock. Further increasing the denaturation strength of the wash buffer had negligible impact on the recovery of poly-ubiquitylated proteins and diGly peptides (R^2^ > 0.99 and 0.93, respectively; Fig. S4B), suggesting that our optimised stringent buffer is sufficient, and also increasing confidence that further non-specifically-bound proteins are only minimally present, if at all.

To evaluate the sensitivity of Ubi-SCAPE, a limited protein input of 100 µg from unstressed cells was tested (Fig. 2B). We identified between 5,411 and 6,753 poly-ubiquitylated protein IDs from this limited basal input, with the equivalent of just 33 µg input equivalent analysed by LC-MS.

To understand further the nature of binding provided by trTUBE-based enrichments, we also evaluated a previously-published, shorter HaloTag–4×trTUBE and a binding-deficient HaloTag– 4×trTUBE, where each ubiquitin-associated domain had been mutated to lower its affinity for ubiquitin (HaloTag–4×trTUBEmut). The 4×trTUBE and 4×trTUBE_mut_ both demonstrated reduced yields, diGly IDs, and reproducibility (Fig. 2B). Notably, the binding-deficient mutant exhibited a uniform, linear ∼25% recovery for poly-ubiquitylated proteins (Fig. S4C), consistent with prior observations;^11^ therefore, we propose that this mutant should be described as “low-binding” rather than “binding-deficient”.

Finally, the stability and batch effects between trTUBE bead preparations were evaluated, by comparing beads stored for 6 months at 4 °C with a freshly-prepared batch (Figs. 2B & S4D). Poly-ubiquitylated protein IDs, diGly IDs, abundances, and reproducibility were almost identical between batches, indicating the robustness of the Ubi-SCAPE tool and method.

### Ubi-SCAPE captures expected poly-ubiquitylome changes with heat-shock

To demonstrate the ability of Ubi-SCAPE to capture biologically-relevant changes in poly-ubiquitylation, we focused on the confident poly-ubiquitylated proteins that were classified as differentially-abundant upon heat-shock, comparing Ubi-SCAPE vs. the standard wash buffer with targeted TEVp-assisted elution (Fig. 3A). Heat-shock is a well-established stress that leads to a global, proteome-wide increase in poly-ubiquitylation.^22, 36^ This directionality of heat-shock-induced changes was far more apparent with our stringent wash conditions, and was almost completely lost with the mild washes at the protein level (Fig. 3A, 2,717 proteins up vs. 553 down with stringent washes, compared with 873 up vs. 747 down with standard washes). More broadly, reduced background contamination from stringent wash conditions substantially improved the quantitative evaluation of the effects of heat-shock, with over 3 times as many poly-ubiquitylated proteins and diGly peptides quantified as increased in abundance in response to stress (2,717 vs. 873 and 981 vs. 326, respectively).

**Figure 3.**
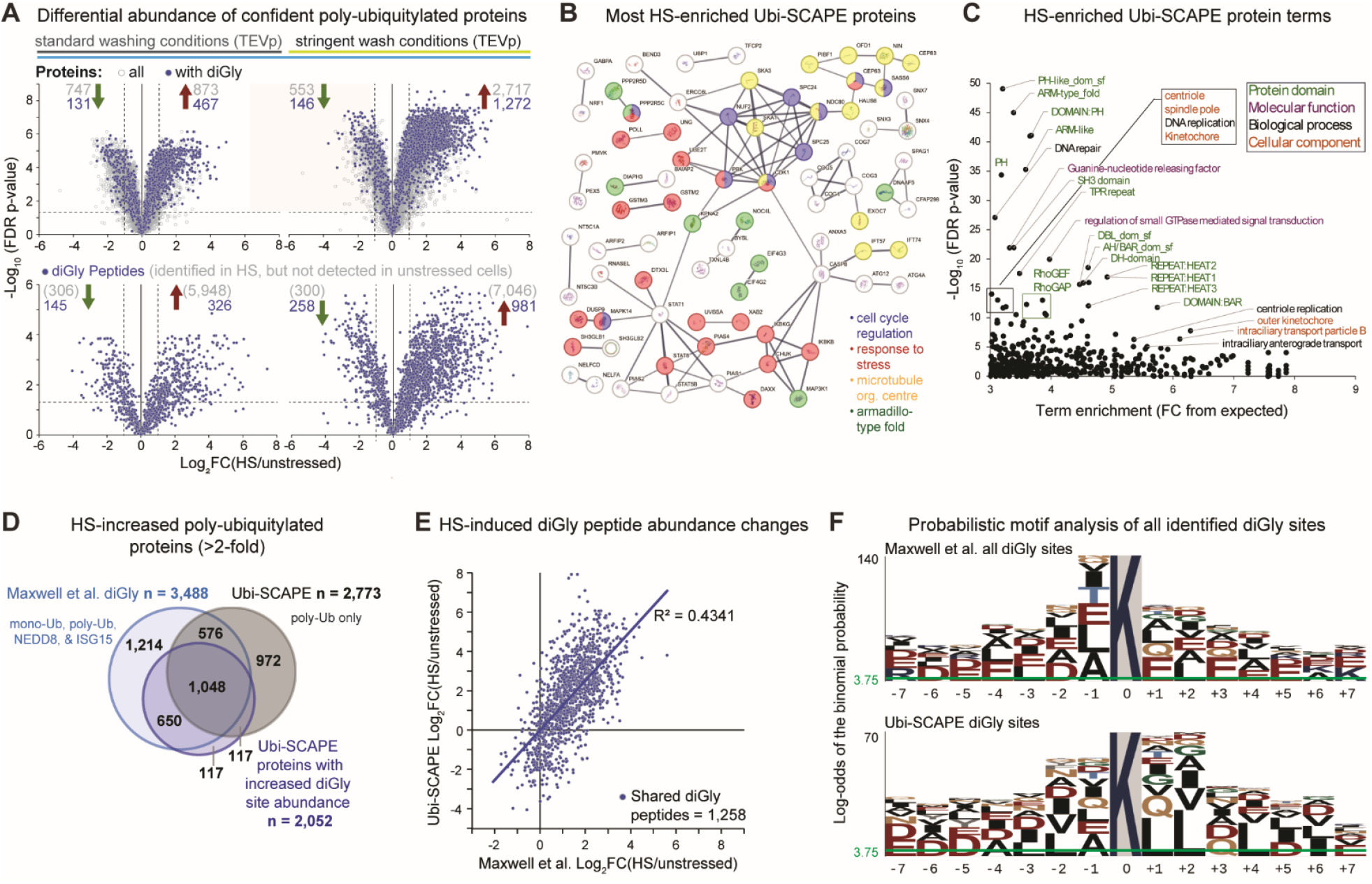
Ubi-SCAPE enables similar biological insights into the heat-shock-induced poly-ubiquitylome as more resource-intensive peptide-level diGly enrichment. (A) Heat-shock-induced changes to abundance of poly-ubiquitylated proteins (with those observed with diGly sites in blue) and diGly peptides, from the experiment described in Fig. 2, with the standard vs. stringent washing conditions. (B) StringDB network of the top 200 significantly over-abundant poly-ubiquitylated proteins with heat-shock, trimmed of unconnected proteins. Confidence > 0.7. (C) Descriptive gene term enrichment for the 2,717 poly-ubiquitylated proteins that increase upon heat-shock. (D) Proteins observed with increased ubiquitylation (and ubiquitin-like modifiers) upon heat-shock by Maxwell et al. using diGly isolation vs. the poly-ubiquitin-specific Ubi-SCAPE. (E) The correlation between diGly peptide abundance changes upon heat-shock identified by Maxwell et al. and Ubi-SCAPE. (F) pLOGO probabilistic motif analysis for sites of poly-ubiquitylation observed by Ubi-SCAPE, alongside a dataset using an anti-diGly antibody to enrich for all diGly peptides (Maxwell et al., 2020), both from heat-shocked HEK293 cells.

Exploration of the gene term enrichment emerging from these 2,717 heat-shock-increased Ubi-SCAPE proteins (Table S4, Fig. S5) identified the well-established signatures of heat-shock (Fig. 3B & 3C).^22, 36^ Furthermore, analysis of enriched protein domains terms indicated several domains that are typically present in tandem repeats (e.g., ARM, HEAT, TPR), which would be more prone to stress-induced aggregation,^37^ and/or feature in molecular chaperones that recognise such heat-aggregating proteins (Fig. 3C).^38, 39^

Finally, we benchmarked our observations with the Maxwell et al. dataset, which used the same cell line and stress (although milder: 43 °C for 1 h).^22^ Their initial ubiquitylomics experiments with a HaloTag–4×TUBE enriched ∼300–1,000 proteins total (depending on the experiment); therefore, they performed their in-depth analyses using diGly-antibody-based peptide isolation. When we compared the diGly peptides captured from our protein-level Ubi-SCAPE workflow with their more resource-intensive diGly-peptide-based enrichment approach, we found that 65% (2,310) of the proteins observed with over two-fold increased ubiquitylation upon heat-shock by diGly peptide isolation were also profiled as such by Ubi-SCAPE, with a further 1,206 unique to Ubi-SCAPE, and 1,214 unique to Maxwell et al.’s diGly dataset (Fig. 3D). Furthermore, for those diGly peptides quantified in both experiments, a correlation (R^2^ = 0.43) and a consistent trend emerged between the two methods (Fig. 3E).

As diGly peptide isolation additionally captures mono-ubiquitylation and some ubiquitin-like modifiers, noteworthy differences were anticipated in the motifs around the ubiquitylated lysine for the diGly peptides identified by the two methods. Surprisingly, there were strong similarities between the motifs, although Ubi-SCAPE exhibited a slight preference for adjacency of the modified lysine to small hydrophobic amino acids (Fig. 3F).

In summary, the Ubi-SCAPE protocol offers high-confidence poly-ubiquitylated protein purification, with low variance, high reproducibility, and high purity, thus providing a robust development towards comprehensive poly-ubiquitylomics.

## CONCLUSIONS

In this study, we introduce a substantial progression in the application of poly-ubiquitin enrichment tools, with Ubi-SCAPE providing greater sensitivity and specificity for the robust and reproducible isolation of over 7,800 poly-ubiquitylated proteins and 8,500 diGly peptides. By combining optimised washing and enzymatic elution, we adapted a streamlined version of trTUBE-based purification for seamless integration into high-throughput and/or high-sensitivity LC-MS proteomics workflows, reducing contamination on non-binding beads to just 2.4% of that observed with the previous workflow. Compared with the more time-, cost-, and input-material-intensive ubiquitin remnant motif antibody-based peptide isolation, Ubi-SCAPE demonstrated similar biological insights, in addition to greater specificity toward poly-ubiquitylation, (i.e., reducing signal from mono- and di-ubiquitylated proteins, and diGly remnants from other ubiquitin-like modifiers such as NEDD8 and ISG15).^6^

Overall, Ubi-SCAPE offers an optimised, streamlined, and highly reproducible protocol for the characterisation of the poly-ubiquitylated proteome to previously unseen depths. The pipeline promises to transform accessibility of the poly-ubiquitylome to researchers, opening new avenues of investigation and understanding of this highly impactful yet understudied PTM.

## Supporting information

Key Resources Table

## ASSOCIATED CONTENT

### Supporting Information

A listing of the contents of each file supplied as Supporting Information are included. The following files are available free of charge:

Figures S1–S5 (PDF)

Tables S1–S4 (XLSX)

Key Resources Table (DOCX)

## AUTHOR INFORMATION

### Author Contributions

The manuscript was written through contributions of all authors. All authors have given approval to the final version of the manuscript.

### Funding Sources

HEJ, GB, SC, and RS were supported by BI’s BBSRC Institute Strategic Programme Grant for Signalling (BB/P013384/1 and BB/Y006925/1); AF and MT by a Wellcome Trust Investigator Award to MT (215542/Z/19/Z). The mass spectrometry analysis was performed at the Newcastle University Laboratory for Biological Mass Spectrometry (the mass spectrometer used in this work was funded by a Wellcome Trust multi-user equipment grant (212947/Z/18/Z)) and Babraham Institute Proteomics Facility funded by BI’s BBSRC Core Capability Grant (BB/CCG2310/1). This work was also supported by a BBSRC Institute Development Grant (BB/IDG2310/1), and through the BBSRC Flexible Talent Mobility Account (FTMA) under grant BB/Z515164/1.

## ACKNOWLEDGEMENTS

The authors would like to thank Yasushi Saeki for the gift of the pRSET–6×TR-TUBE (Addgene plasmid #110313); MRC PPU Reagents and Services facility for the pET28a–HALO–TEV (#DU23222); Daniel Daley for pET28a T7pCONS TIR-2 sfGFP (Addgene #306150); Yimon Aye for pET28a His6-HaloTag-Nrf2 (Addgene #62455); Sam Lord and Yu-Chiang Lai for plasmids containing 4×trTUBE (#DU58810) and 4×trTUBE_mut_ (#DU58829), generated by MRC PPU Reagents and Services facility; and Shannon Blay and Madel Tutor for cloning of 6×His tag into 6×His-HaloTag-TEVc-6×trTUBE. Babraham Institute Proteomics Facility performed LC-MS analysis of samples during method development. The authors declare no competing financial interests.

## Abbreviations

diGly: Lys-ε-GlyGly ubiquitin tryptic remnant motif
IP: immunoprecipitation
polyUb: poly-ubiquitin
PTM: post-translational modification
rpm: revolutions per minute
SDC: sodium deoxycholate
SDS: sodium dodecyl sulfate
SDS-PAGE: sodium dodecyl sulfate-based polyacrylamide gel electrophoresis
SPD: samples-per-day
TEVp: Tobacco Etch Virus protease
TEVc: Tobacco Etch Virus protease cleavage sequence (ENFLQG)
TFA: trifluoroacetic acid
trTUBE: trypsin-resistant tandem ubiquitin binding entity
Ub: ubiquitin
Ubi-SCAPE: Ubiquitylomics by Stringent, Cleavable, Affinity-based Proteome Extraction

## Notes

### Competing Interest Statement

The authors have declared no competing interest.

