## Supplementary Information for "Ubi-SCAPE enables deep exploration of the poly-ubiquitylome"

### Supplementary Figures

**Figure S1.** Evaluation of magnetic HaloTag bead saturation with *E. coli* lysate expressing HaloTag–TEVc–6×trTUBE.

**Figure. S2**. Initial evaluation of tolerated binding conditions for poly-ubiquitin chains with HaloTag–TEVc–6×trTUBE.

**Figure S3.** Further evaluation of binding conditions of HaloTag–TEVc–6×trTUBE to varying lengths of poly-ubiquitin chains.

**Figure S4.** Proteomics evaluation of trTUBE variants and buffers.

**Figure S5**. Top 6 enriched terms per DAVID category among the Ubi-SCAPE heat-shock-enriched proteome.

### ADDITIONAL INFORMATION

**Table S1-S4** (separate .xlsx spreadsheet):

**Table S1** The protein group IDs for the trTUBE and control experiments

**Table S2** Ubiquitin remnant motif peptide IDs

**Table S3** MS2-diaPASEF IM-m/z window definition

**Table S4** DAVID–generated terms list for proteins with significantly increased poly-ubiquitylation upon heat shock

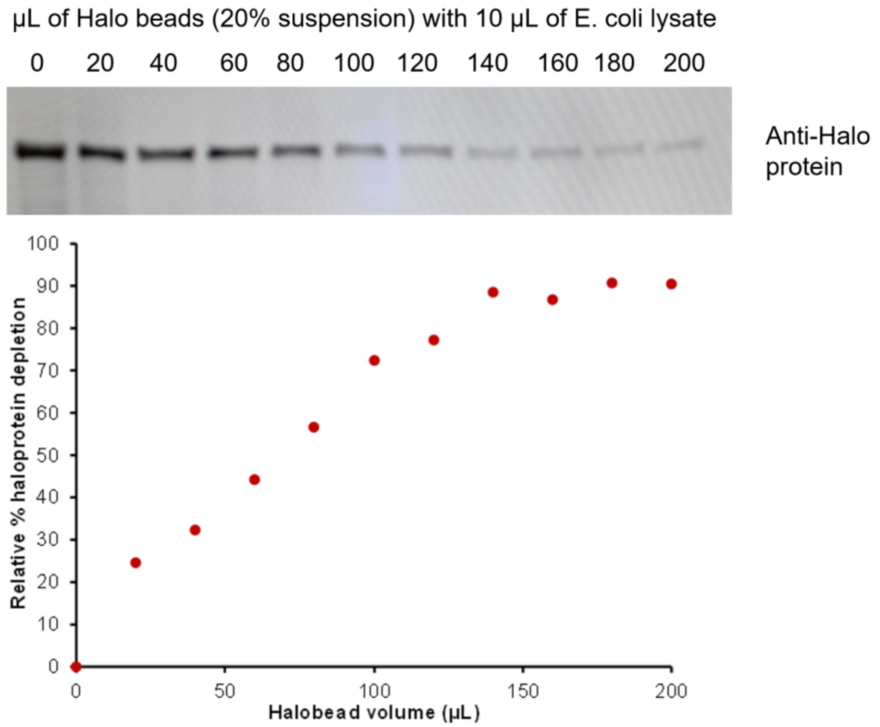


**Figure S1. Evaluation of magnetic HaloTag bead saturation with *E. coli* lysate expressing HaloTag–TEVc–6×trTUBE.** Increasing volumes of Magne-HaloTag beads were added to 10 µL of *E. coli* lysate (prepared as a 3:1 ratio of lysis buffer:pellet volume), followed by incubation for 2 h at room temperature in a 100 µL binding reaction on a thermomixer at 1,000 rpm with intermittent mixing (3 s on, 3 s off). The supernatant was subsequently collected and 18 µL evaluated by SDS–PAGE and immunoblotting with a mouse anti-HaloTag antibody. Beads appeared fully saturated at a beads:lysate ratio of 14:1, where no further bead addition could deplete further HaloTag protein. To accommodate for batch effects, this was rounded up to a ratio of 200 µL lysate per 1 mL Magne-HaloTag bead slurry for all experiments.

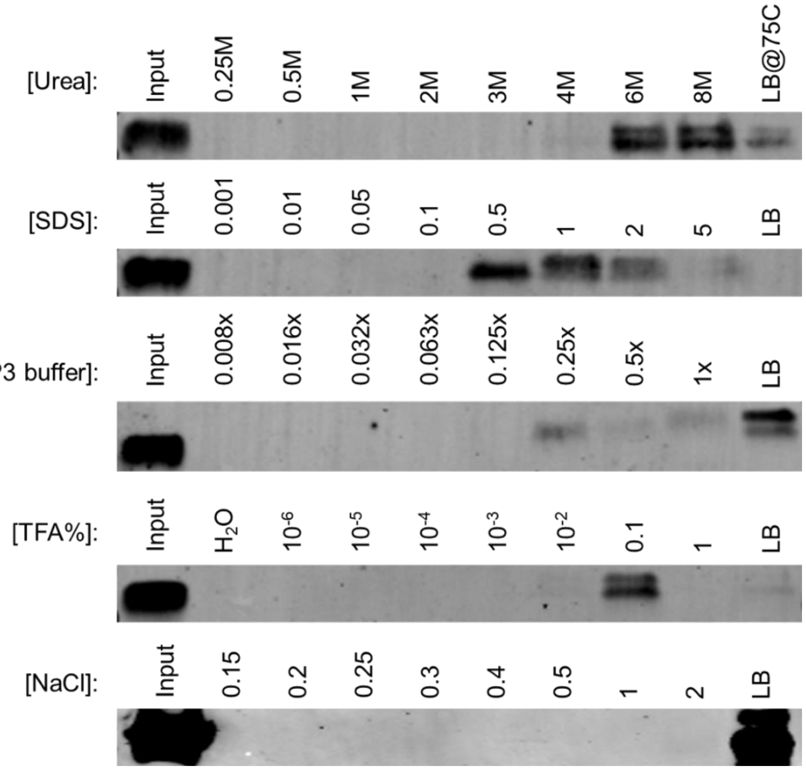


**Figure S2. Initial evaluation of tolerated binding conditions for poly-ubiquitin chains with HaloTag–TEVc–6×trTUBE.** 10 µL equivalent of Magne-HaloTag beads were incubated with a 100 ng mixture of seven tetra-ubiquitin chains (M1, K6, K11, K29, K33, K48, and K63) in 1 mL IP buffer (50 mM Tris, 150 mM NaCl, 1% IGEPAL CA-630) with 0.1 mg/mL BSA overnight at 4 °C with end-over-end rotation. Beads were washed three times in ice-cold IP buffer and subjected to 15 µL volumes of the buffers indicated in each lane sequentially, with each supernatant collected for evaluation by SDS–PAGE and anti-ubiquitin (VU-1) immunoblotting. Each reaction was conducted on a thermomixer for 2 min at 4 °C, 800 rpm. Urea, SDS, NaCl, & SP3 buffer were diluted with IP buffer, whereas TFA was serially diluted with water. A final elution using loading buffer and heat-based elution at 75 °C for 10 min at 800 rpm was used to quantify remaining bead-bound poly-ubiquitin. This experiment established the strongest tolerable trTUBE:poly-ubiquitin binding and wash conditions to help eliminate non-specific background protein binding.

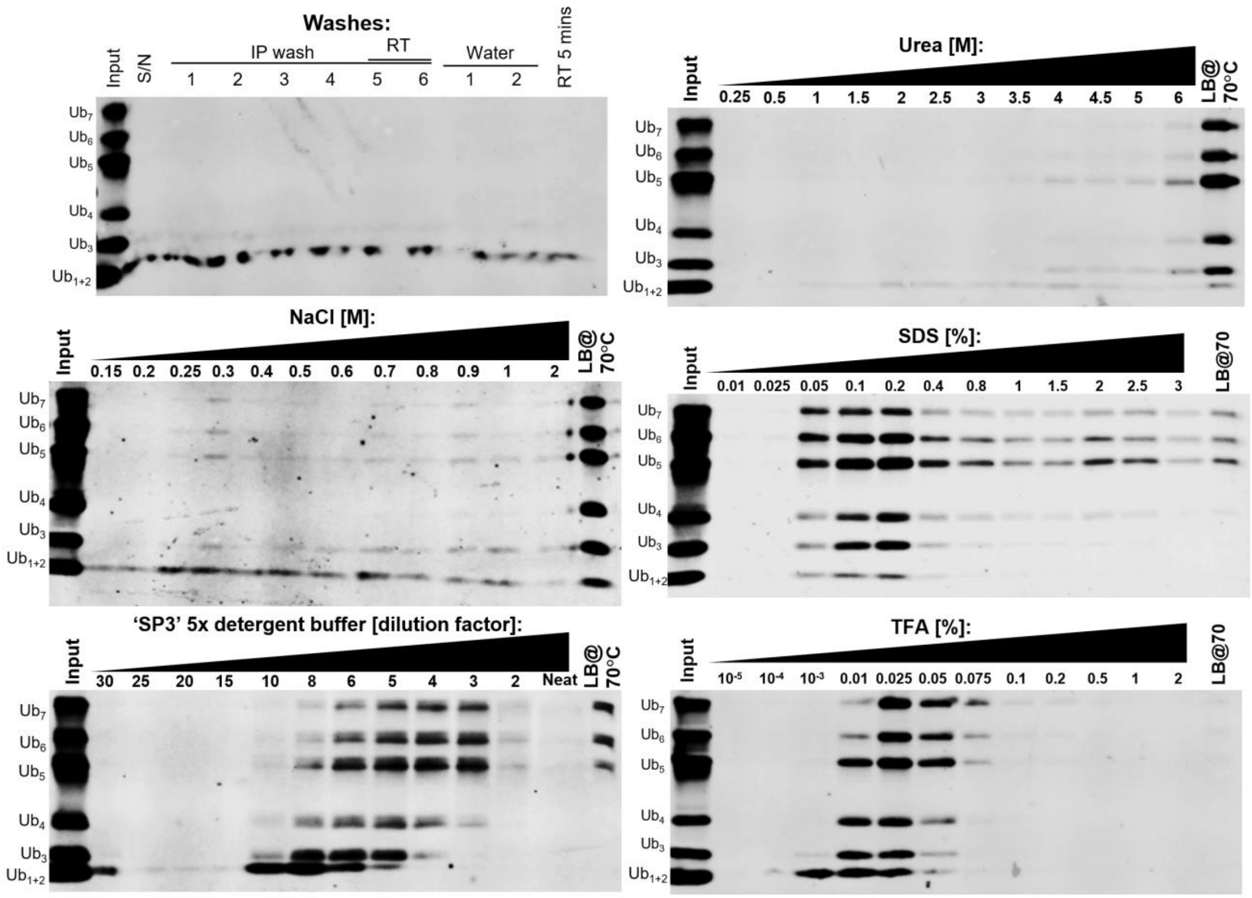


**Figure S3. Further evaluation of binding conditions of HaloTag–TEVc–6×trTUBE to varying lengths of poly-ubiquitin chains.** Experiment performed as in Fig. S2, except with 1.2 μg of linear (M1-linked) poly-ubiquitin chains incubated with HaloTag–TEVc–6×trTUBE per serial evaluation. The top left blot details initial washes of all 5 sets of beads in IP buffer at 4 °C or room temperature to check temperature stability and effects of washes with ultrapure water (to ensure binding was not affected by osmolarity). Mono- and di-ubiquitin were observed with elution with all tested parameters, demonstrating the reduced stability of their interaction (presumably due to avidity effects).

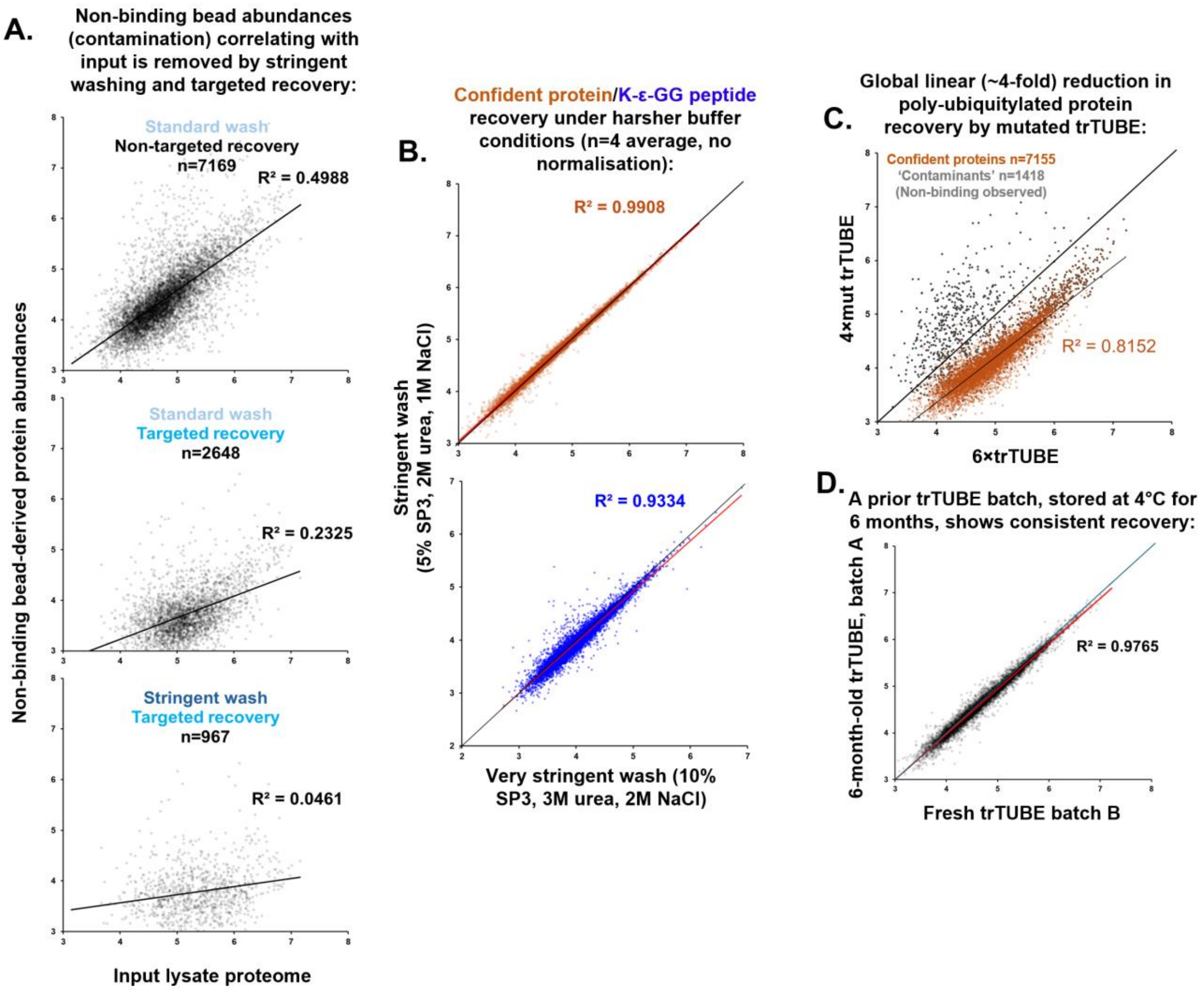


**Figure S4. Proteomics evaluation of HaloTag–trTUBE variants and buffers.** Based on the data presented in Fig. 2B. **A.** The Ubi-SCAPE protocol eliminates any correlation with the input proteome for non-specific binding controls. Pearson correlation coefficients ($R^2$) between HaloTag–TEVc –empty non-binding-derived contaminants and the input proteome for the three compared method variations (on-bead digestion with standard washing, TEVp-based elution and standard washing, and TEVp-based elution with optimised washing) are shown. The combination of these two modifications to the protocol eliminated almost all correlation among the residual observations. **B.** Ubi-SCAPE wash buffer is sufficient to remove contaminants. The abundances of confident pUb-proteins and diGly peptides when using trTUBE isolations processed using an even more stringent wash buffer (50 mM Tris pH 7.4, 1% IGEPAL, 2 M NaCl, 3 M urea, 0.1% SDS, 0.1% SDC, 0.1% Tween, 0.1% Triton-X)) versus that described as the standard method (50 mM Tris pH 7.4, 1% IGEPAL, 1 M NaCl, 2 M urea, 0.05% SDS, 0.05% SDC, 0.05% Tween, 0.05% Triton-X) to understand if a more stringent version of the method was possible. Given negligible differences were observed between the two wash buffers, it was concluded that either buffer was sufficient to remove contaminants, and further benefits from stronger washing were unlikely and based on Figure S2–S4 would risk substantial losses. **C.** Mutant trTUBE binds poly-ubiquitylated proteins with ~25% recovery vs. 6×trTUBE. Protein abundance was compared between beads saturated with HaloTag–TEVc–6×trTUBE and HaloTag–TEVc–4×trTUBE$_{mut}$. The consistent linear reduction in polyUb-protein binding indicated a reduced but non-negligible binding capacity. **D.** Ubi-SCAPE is highly reproducible even between bead batches. A stability and batch-effect control for HaloTag–TEVc–6×trTUBE-bound beads stored for 6 months at 4 °C was compared with a freshly-prepared batch with the same lysate.

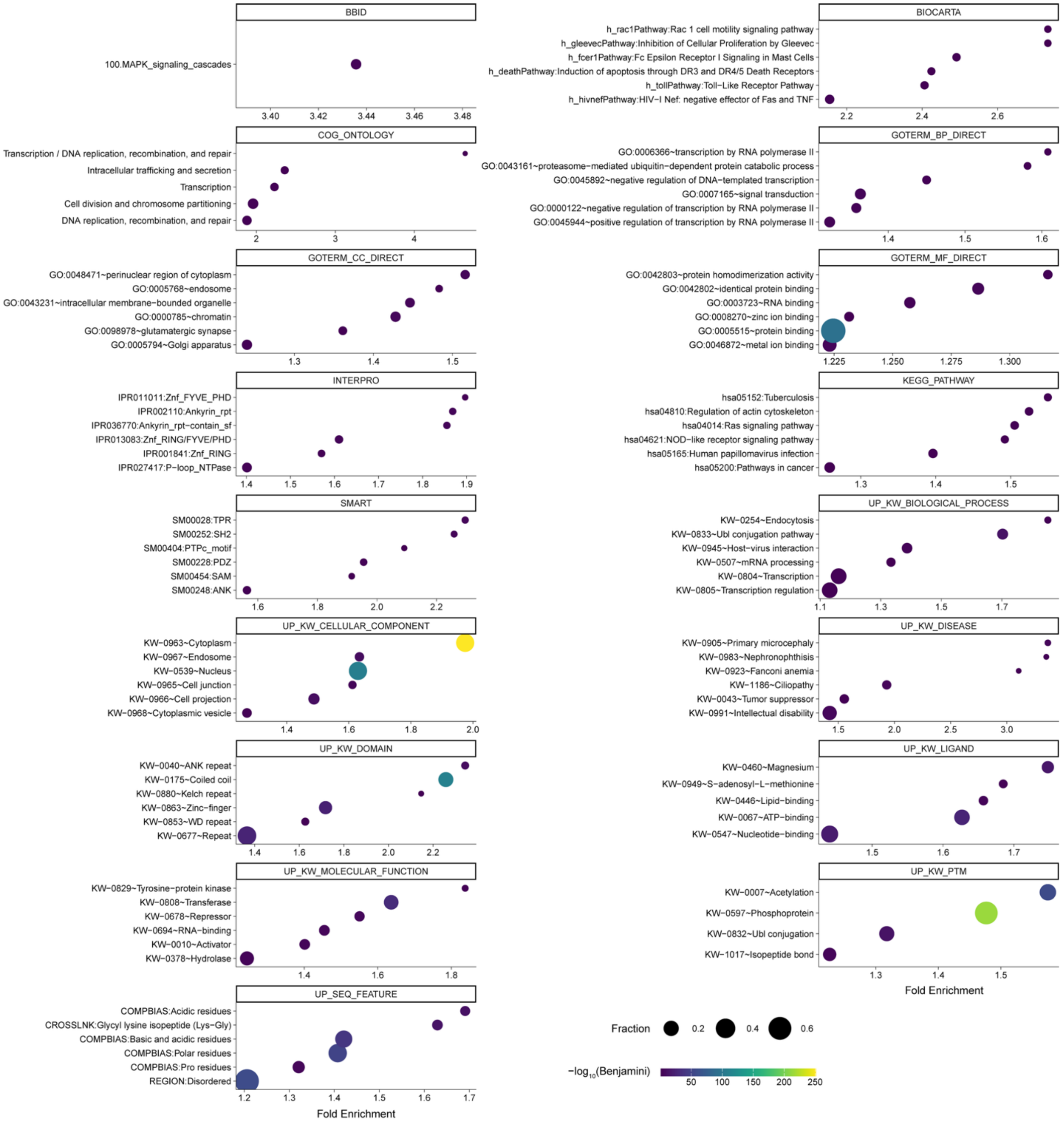


**Figure S5. Top 6 enriched terms per DAVID category among the Ubi-SCAPE heat-shock-enriched proteome.** Data from Supplementary Table S4. Only terms with a Fold Enrichment > 1.0 and Benjamini-Hochberg adjusted p-value < 0.05 are plotted. 'Fraction' indicates the proportion of the whole term represented by the enriched proteins (i.e., 'Pop Hits' divided by 'Pop Total' in Supplementary Table S4).
