## Supplementary material for "Ubi-SCAPE enables deep exploration of the poly-ubiquitylome": Key Resources Table

| **REAGENT or RESOURCE** | **SOURCE** | **IDENTIFIER** |
| --- | --- | --- |
| **Antibodies** | | |
| Anti-Ubiquitin, mouse monoclonal | LifeSensors | Cat# VU101; RRID:AB_2716558 |
| Anti-HaloTag, mouse monoclonal | Promega | Cat# G9211;  RRID:AB_2688011 |
| Anti-Mouse IgG (H+L), DyLight 680, goat | Cell Signaling Technology | Cat# 5470; RRID:AB_10696895 |
| Anti-Mouse IgG (H+L), DyLight 800, goat | Cell Signaling Technology | Cat #5257; RRID:AB_10693543 |
| **Bacterial and virus strains** |  |  |
| *E. coli:* DH5-alpha (sub-cloning efficiency) | Thermo | 18265017 |
| *E. coli:* Rosetta 2(DE3) (Novagen) | Sigma | 71397 |
| **Chemicals, peptides, and recombinant proteins** | | |
| DMEM, high glucose | Gibco | 41965039 |
| L-Glutamine (200 mM) | Gibco | 25030081 |
| Penicillin-Streptomycin (10,000 U/mL) | Gibco | 15140122 |
| MEM Non-Essential Amino Acids Solution (100X) | Gibco | 11140050 |
| HyClone Characterized Fetal Bovine Serum, U.S. Origin, Heat-Inactivated | Cytiva | SH30071.02HI |
| Trypsin-EDTA (0.05%), phenol red | Gibco | 25300054 |
| Isopropyl beta-D-1-thiogalactopyranoside (IPTG) | Melford Laboratories | I56000 |
| Sodium chloride | VWR | X190 |
| Sodium hydroxide | Fisher | S/4920/53 |
| Phosphate buffered saline (PBS) | Sigma | P2272 |
| Pierce Universal Nucleases for Cell Lysis | Thermo | 88701 |
| HEPES (N-(2-Hydroxyethyl) piperazine N'-(2-ethanesulfonic acid)) | Melford Laboratories | H75030 |
| Ammonium bicarbonate | Sigma | A6141 |
| Urea | VWR | 28877 |
| Triton X-100 | Sigma | T9284 |
| IGEPAL CA‑630 | Sigma | 56741 |
| Tween 20 | Sigma | P1379 |
| Sodium dodecyl sulfate (SDS) | Bio-Rad | 161-0301 |
| Sodium deoxycholate | Sigma | D6750 |
| Tris-(2-carboxyethyl)phosphine (TCEP), hydrochloride | Thermo | T2556 |
| cOmplete mini protease inhibitor tablet, EDTA-free (Roche) | Sigma | 1836170 |
| Phenylmethanesulfonyl fluoride (PMSF) | Sigma | P7626 |
| 2-chloroacetamide | Sigma | C0267 |
| PR-619 | Sigma | 662141 |
| Orange G, Electrophoresis Grade | Thermo | J62743 |
| Trypsin/Lys-C Mix, Mass Spec Grade | Promega | V5073 |
| Acetonitrile (ACN), LC-MS-grade | Fisher | A9551 |
| Formic Acid, LC-MS-grade | Thermo | A11750 |
| Water, HPLC-grade | Fisher | W/0110/PB17 |
| TMTpro 18-plex Isobaric Label Reagent | Thermo | A52045 |
| Triethylammonium bicarbonate (TEAB) | Thermo | 90114 |
| Hydroxylamine | Thermo | 90115 |
| Mono-ubiquitin, human (recombinant) | Sigma | U5507 |
| Poly-ubiquitin chains (Ub2-7), (linear) (recombinant) | Enzo | BML-UW1010 |
| K6-linked di-ubiquitin, human (recombinant) | South Bay Bio | SBB-UP0060 |
| K11-linked di-ubiquitin, human (recombinant) | South Bay Bio | SBB-UP0063 |
| K29-linked di-ubiquitin, human (recombinant) | South Bay Bio | SBB-UP0077 |
| K33-linked di-ubiquitin, human (recombinant) | South Bay Bio | SBB-UP0066 |
| K48-linked di-ubiquitin, human (recombinant) | South Bay Bio | SBB-UP0069 |
| K63-linked di-ubiquitin, human (recombinant) | South Bay Bio | SBB-UP0072 |
| K6-linked tetra-ubiquitin, human (recombinant) | South Bay Bio | SBB-UP0061 |
| K11-linked tetra-ubiquitin, human (recombinant) | South Bay Bio | SBB-UP0064 |
| K29-linked tetra-ubiquitin, human (recombinant) | South Bay Bio | SBB-UP0078 |
| K33-linked tetra-ubiquitin, human (recombinant) | South Bay Bio | SBB-UP0067 |
| K48-linked tetra-ubiquitin, human (recombinant) | South Bay Bio | SBB-UP0070 |
| K63-linked tetra-ubiquitin, human (recombinant) | South Bay Bio | SBB-UP0073 |
| **Critical commercial assays** | | |
| Monarch Spin DNA Gel Purification Kit | New England Biolabs | T1120S |
| Monarch Spin Plasmid Miniprep Kit | New England Biolabs | T1110 |
| Pierce BCA Protein Assay Kit | Thermo | 23225 |
| Pierce Quantitative Peptide Assay - Colorimetric | Thermo | 23275 |
| **Deposited data** | | |
| Various TUBEs and conditions incubated with HEK293 +/- heat-shock lysates (Fig. 2) | This study | ProteomeXchange: PXD073597 |
| **Experimental models: Cell lines** | | |
| HEK293 | ATCC | Cat# CRL-1573; RRID:CVCL_0045 |
| **Oligonucleotides (all sequences 5’ to 3’)** | | |
| Cloning PCR primer 108-BlpI-F1: ATTATGCTTAGCGGCAGCAGCCATC | This study |  |
| Cloning PCR primer pET28a-BlpI-R1: TGCTAGTTATTGCTCAGCGGTGGCA | This study |  |
| **Recombinant DNA** | | |
| pET28a−HaloTag−TEVc−[MCS] | MRC-PPU Reagents & Services, UK | Cat# DU23222 |
| pET28a T7pCONS TIR-2 sfGFP | Daniel Daley, (Addgene plasmid #154464; <http://n2t.net/addgene:154464>) | RRID: Addgene_154464 |
| pRSET−6×TR-TUBE | Yasushi Saeki (Addgene plasmid #110313; <http://n2t.net/addgene:110313>) | RRID: Addgene_110313 |
| pET28a HaloTag−TEVc−empty | This study | Internal# pRSS113 |
| pET28a HaloTag−TEVc−6×trTUBE | This study | Internal# pRSS114 |
| pET28a 6×His−TEVc−HaloTag−4×trTUBE | Yu-Chiang Lai, U. Birmingham, UK & MRC-PPU Reagents & Services, UK | Cat# DU58810 |
| pET28a 6×His−TEVc−HaloTag−4×trTUBE_mut_ | Yu-Chiang Lai, U. Birmingham, UK & MRC-PPU Reagents & Services, UK | Cat# DU58829 |
| pET28a 6×His−HaloTag−Nrf2 | Yimon Aye (Addgene plasmid #62455; <http://n2t.net/addgene:62455>) | RRID: Addgene_62455 |
| pET28a 6×His−HaloTag−TEVc−6×trTUBE | This study | Internal# pRSS339 |
| pET28a 6×His−HaloTag−TEVc−4×trTUBE | This study | Internal# pRSS342 |
| pET28a 6×His−HaloTag−TEVc−4×trTUBE_mut_ | This study | Internal# pRSS343 |
| **Software and algorithms** | | |
| DIA-NN (v1.9) | <https://github.com/vdemichev/DiaNN> | Demichev et al. (2020) <https://doi.org/10.1038/s41592-019-0638-x> |
| R (v4.2.2) | <https://cran.r-project.org/> |  |
| diann-rpackage (v1.9) | <https://github.com/vdemichev/diann-rpackage> | Demichev et al. (2020) <https://doi.org/10.1038/s41592-019-0638-x> |
| DAVID: Database for Annotation, Visualization, and Integrated Discovery (v2023q4) | <https://davidbioinformatics.nih.gov/> | Sherman et al. (2022) <https://doi.org/10.1093/nar/gkac194> |
| STRING protein-protein interaction database (v11) | <https://string-db.org/> | Szklarczyk et al. (2023) <https://doi.org/10.1093/nar/gkac1000> |
| **Other** | | |
| timsTOF HT | Bruker | <https://www.bruker.com/en/products-and-solutions/mass-spectrometry/timstof/timstof-ht.html> |
| Orbitrap Eclipse Tribrid Mass Spectrometer | Thermo | <https://www.thermofisher.com/order/catalog/product/FSN04-10000> |
| LiCor Odyssey CLx Imager | LI-COR Biotech | <https://www.licorbio.com/support/answer-portal/imaging-systems/odyssey-clx.html> |
| Savant SpeedVac SPD210 Vacuum Concentrator | Thermo | SPD210P2-230 |
| Thermomixer comfort 5355 | Eppendorf | <https://www.eppendorf.com/product-media/doc/en/085934_Operating-Manual/Eppendorf_Sample-Preparation_Operating-manual_Thermomixer-comfort_Thermomixer-R.pdf> |
| 0.5 mL Protein LoBind Tube | Eppendorf | 0030108094 |
| 2 mL Protein LoBind Tube | Eppendorf | 0030108132 |
| Magne-HaloTag Beads | Promega | G7281 |

**Buffers**

| **BUFFER** | **COMPOSITION** |
| --- | --- |
| Bacterial Lysis Buffer | 50 mM Tris‑HCl pH 7.5, 150 mM NaCl, 1% Triton X‑100, 20 mM TCEP, 1× cOmplete protease inhibitors, 0.2 mM PMSF |
| IP Buffer | 50 mM Tris-HCl pH 7.5, 150 mM NaCl, 1% IGEPAL CA‑630 |
| Urea SP3 Lysis Buffer | 50 mM HEPES pH 8.0, 8 M urea, 1% SDS, 1% Triton X‑100, 1% IGEPAL CA-630, 1% Tween 20, 1% sodium deoxycholate, 150 mM NaCl, 10 mM TCEP, 1x cOmplete protease inhibitors, EDTA-free, 1 mM PMSF, 10 µM PR-619, 40 mM 2‑chloroacetamide (CAA), 20 U/μL Pierce Universal Nucleases |
| IP500 Wash Buffer | IP Buffer adjusted to 500 mM NaCl |
| stringent trTUBE Wash Buffer | IP Buffer supplemented with 5% SP3 buffer, 2 M urea, 1 M NaCl |
| very stringent trTUBE Wash Buffer | IP Buffer supplemented with 10% SP3 buffer, 3 M urea, 2 M NaCl |
| SDS–PAGE Loading Buffer | 25 mM Tris-HCl, pH 6.8, 10% glycerol, 1% SDS, 0.05% (w/v) Orange G |
